# Differential Expression of Coding and Long Noncoding RNAs in Keratoconus-affected Corneas

**DOI:** 10.1101/267054

**Authors:** Mariam Lofty Khaled, Yelena Bykhovskaya, Sarah E.R. Yablonski, Hanzhou Li, Michelle D. Drewry, Inas F. Aboobakar, Amy Estes, Xiaoyi Gao, W. Daniel Stamer, Hongyan Xu, R. Rand Allingham, Michael A. Hauser, Yaron S. Rabinowitz, Yutao Liu

## Abstract

**PURPOSE:** Keratoconus (KC) is the most common corneal ectasia. We aimed to determine the differential expression of coding and long noncoding RNAs (lncRNAs) in human corneas affected with KC.

**METHODS:** 200ng total RNA from the corneas of 10 KC patients and 8 non-KC normal controls was used to prepare sequencing libraries with the SMARTer Stranded RNA-Seq kit after ribosomal RNA depletion. Paired-end 50bp sequences were generated using Illumina HiSeq 2500 Sequencer. Differential analysis was done using TopHat and Cufflinks with a gene file from Ensembl and a lncRNA file from NONCODE. Pathway analysis was performed using WebGestalt. Using the expression level of differentially expressed coding and noncoding RNAs in each sample, we correlated their expression levels in KC and controls separately and identified significantly different correlations in KC against controls followed by visualization using Cytoscape.

**RESULTS:** Using |fold change| ≥ 2 and a false discovery rate ≤ 0.05, we identified 436 coding RNAs and 584 lncRNAs with differential expression in the KC-affected corneas. Pathway analysis indicated the enrichment of genes involved in extracellular matrix, protein binding, glycosaminoglycan binding, and cell migration. Our correlation analysis identified 296 pairs of significant KC-specific correlations containing 117 coding genes enriched in functions related with cell migration/motility, extracellular space, cytokine response, and cell adhesion, suggesting the potential functions of these correlated lncRNAs, especially those with multiple pairs of correlations.

**CONCLUSIONS:** Our RNA-Seq based differential expression and correlation analyses have identified many potential KC contributing coding and noncoding RNAs.

## Introduction

Keratoconus (KC) is the most common corneal ectatic disorder and a leading indicator for corneal transplantation in developed countries^1,2^. The cornea is the outermost, avascular, transparent part of the eye and maintenance of its shape and transparency is critical for optimizing the eye’s refractive power^3^. KC is characterized by bulging and thinning cone-shaped cornea, resulting in myopia, irregular astigmatism, and, eventually, visual impairment. KC usually starts at puberty and progresses until it stabilizes in the third or fourth decade^4,5^. KC is a complex multifactorial disorder, and changes in numerous genes and environmental factors are thought to be responsible for the disease development and progression^6^. KC patients report significantly impaired vision-related quality of life which worsens with time due to disease progression^7^. Visual impairment develops from myopia and irregular astigmatism^8^.

Next-generation sequencing is currently used as a tool for high-throughput assessment of nucleotide sequences that is amenable to various applications including whole genome, exome, and RNA sequencing (RNA-Seq). RNA-Seq allows the entire transcriptome to be surveyed in a high-throughput and quantitative manner^9,10^. Consequently, RNA-Seq has been used to identify various populations of RNA, including total RNA, pre-mRNAs, and noncoding RNAs (ncRNAs), such as microRNA, snoRNAs, and long ncRNAs (lncRNAs)^11,12^ LncRNAs are class of noncoding RNAs that are at least 200 nucleotides long^13,14^. LncRNAs have been found to regulate gene expression transcriptionally and post-transcriptionally in both physiological and pathological conditions ^14,15^.

Previous studies have started the screening process attempting to understand processes that underlie KC using different approaches. Wentz-Hunter et al. have performed targeted expression analysis of the stromal layer of the cornea to identify the differentially expressed transcripts in KC versus normal cornea^16^. Soon afterwards, microarray analysis was used to assess RNAs isolated from either the KC-affected epithelium or keratocytes ^17,18^. By 2005, differentially expressed proteins in KC patients have been investigated using either the whole cornea, certain layer, or tears^19–22^. In 2017, Szcześniak et al. and Kabza M et al. used RNA-Seq to determine differentially expressed RNAs between KC-affected and other disease affected corneas ^23^.

In this study for the first time we used high-throughput RNA-Seq to identify the differentially expressed coding and lncRNAs in KC-affected corneas versus non-diseased normal ones. This is the first report to study the correlation between coding RNAs and lncRNAs, and the impact of these differentially expressed functioning RNAs on the pathogenesis of KC. Additionally, we used the droplet digital PCR (ddPCR) technique to validate the differential expression of selected coding RNAs and lncRNAs which are potentially related to the disease pathogenesis. The identified differentially expressed RNAs may provide a better understanding of the functions of lncRNAs and the regulatory signaling pathways involved in KC pathogenesis.

## Methods

### Human Cornea Samples and RNA Extraction

We followed the Tenets of Declaration of Helsinki. Unaffected human cornea samples were collected from postmortem donors from North Carolina Eye Bank, and the KC-affected corneal samples were collected during corneal transplantation surgery at the Cedars-Sinai Medical Center. Both studies have been approved by the Institutional Research Board offices at Duke University School of Medicine and Cedars-Sinai Medical Center, respectively. Written informed consent was obtained from all the patients at the Cedars-Sinai Medical Center. This study included ten KC-affected corneas and 8 normal non-KC corneas (**Table 1**). The inclusion and exclusion criteria of KC diagnosis has been described previously ^24,25^. The postmortem donors had no other ocular disorders affecting their cornea. The cornea was stored in the RNALater™ RNA stabilization reagent (Qiagen, Valencia, CA, USA) to preserve RNA after collection. Total RNA was extracted from the whole cornea tissue using mirVana miRNA isolation kit from ThermoFisher Scientific, and with NucleoSpin^®^ RNA/Protein isolation kit (MACHEREY-NAGEL Inc., Bethlehem, PA), according to the manufacturer’s recommendations. RNA quality was evaluated using Agilent 2100 Bioanalyzer with Agilent’s RNA 6000 Pico Kit (Agilent, Santa Clara, CA, US). We selected RNA samples with a minimum RIN (RNA Integrity Number) score of 6 for RNA-Seq.

**Table 1.** Clinical phenotypes of patients with KC and normal controls.

| **Sample** | **Age (years)** | **Race** | **Sex** |
| --- | --- | --- | --- |
| KC1 | 34 | Hispanic | Male |
| KC2 | 42 | Caucasian | Male |
| KC3 | 35 | Caucasian | Female |
| KC4 | 41 | Caucasian | Male |
| KC5 | 43 | Caucasian | Female |
| KC6 | 50 | Caucasian | Male |
| KC7 | 64 | Caucasian | Female |
| KC8 | 23 | Hispanic | Male |
| KC9 | 42 | Caucasian | Male |
| KC10 | 24 | Hispanic | Male |
| Control1 | 66 | Caucasian | Female |
| Control2 | 67 | Caucasian | Male |
| Control3 | 61 | African | Female |
| Control4 | 76 | Caucasian | Female |
| Control5 | 77 | Caucasian | Male |
| Control6 | 49 | African-American | Male |
| Control7 | 66 | Caucasian | Male |
| Control8 | 73 | Caucasian | Male |

### Stranded Total RNA-Seq

We used 200 ng high-quality total RNA from the corneas of 10 KC patients and 8 normal controls to prepare sequencing libraries with the TaKaRa SMARTer Stranded RNA-Seq kit after ribosomal RNA depletion using the RiboGone - Mammalian kit from TaKaRa (TaKaRa Bio USA, Inc., Mountain View, CA, USA). Briefly, the RiboGone – Mammalian kit removes ribosomal RNA (rRNA) and mitochondrial RNA (mtRNA) sequences from human total RNA samples based on hybridization and RNase H digestion that specifically depletes 5S, 5.8S, 18S, and 28S nuclear rRNA sequences, as well as 12S mtRNA sequences. The rRNA-depleted total RNA was reverse-transcribed to synthesize the first-strand cDNA with SMARTer Stranded oligo at the 5’-end and tailed with SMART Stranded N6 primer at the 3’-end. The purified first-strand cDNA was amplified by PCR using universal forward PCR primer and reverse PCR indexing primer set. RNA-Seq library was purified using SPRI AMPure Beads (Beckman Coulter, Inc., Indianapolis, IN, USA) and validated using the Agilent 2100 Bioanalyzer with Agilent’s High Sensitivity DNA Kit (Agilent, Santa Clara, CA, USA). Pooled RNA-Seq libraries were loaded into an Illumina HiSeq 2500 sequencer (Illumina, Inc., San Diego, CA, USA) and sequenced with paired-end 50bp reading using the rapid mode at the Georgia Cancer Center Integrated Genomics Core of Augusta University (http://www.augusta.edu/cancer/research/shared/genomics/index.php).

After quality check and control with all the sequencing reads, demultiplexed reads were aligned by TopHat using paired-end reading with the approximation of the median library size. Counts of sequencing reads were normalized using Cufflinks in fragments per kilo bases and millions reads (FPKM). After normalizing the sequencing read counts, we annotated the transcriptswith a coding gene file from Ensembl database at the gene isoform level. We performed differential expression analysis using the Cuffdiff package^26^. We used a fold change of at least 2 and false discovery rate (FDR) ≤0.05 to identify genes with significant expression changes in KC corneas versus normal ones. The list of differentially expressed coding genes was loaded to the Web-based Gene Set Analysis Tookit (WebGestalt) (www.webgestalt.org) 2013 version for gene ontology enrichment and KEGG pathway analysis^27^. For lncRNAs, we annotated them using a file from the NONCODE version 5 database^28^. After normalization using Cufflinks, we performed differential expression analysis using Cuffdiff. For lncRNA data, missing expression data in specific samples was replaced with a value of 0.001 to enable the differential analysis between cases and controls. For lncRNA differential expression analysis, to avoid the possible impact of potential DNA contamination and RNA isolation bias, we required a minimal size of 300bp for all the lncRNAs. Analysis resulted in a list of lncRNAs with the corresponding chromosome position, fold changes, and adjusted FDR value. We used |fold change| ≥ 2 and FDR ≤ 0.05 to identify lncRNAs with significant expression changes in KC corneas vs. normal ones.

### Validation using droplet digital PCR technique (ddPCR)

Twenty-one differentially expressed coding RNAs were chosen for validation through quantitative droplet digital PCR technique (ddPCR) as previously described. We selected these coding RNAs based on the following criteria: 1. reasonably high corneal expression; 2. relevance to corneal integrity, transparency, and cellular adhesion; 3. potential involvement in oxidative stress; and 4. related with corneal wound healing. Specific EvaGreen-based ddPCR assays for these 21 genes and 4 the reference genes (*GAPDH*, *PPIA*, *HPRT1*, and *RPLP0*) were obtained from Bio-Rad Laboratories, Inc. (Hercules, CAUSA). To validate the differentially expressed lncRNAs, we selected 7 lncRNAs which have reasonable corneal expression and strong correlations with various other coding and lncRNAs. Specific ddPCR expression assays with specific probes were designed and ordered using the online ddPCR assay design tool with the specific RNA sequence of each selected lncRNA for copy number determination (https://www.bio-rad.com/digital-assays/#/). We used RNA samples from the total RNA-Seq experiment for validation: 8 KC-affected corneas and 8 control post mortem corneas. RNA utilized for ddPCR was reverse transcribed using Superscript IV (Invitrogen, Thermo Fisher Scientific, ILL, USA) with random hexamers that anneal to random complementary sites on a target RNA, following the manufacturer protocol. The generated cDNA samples were further diluted 5-fold using ultrapure DNase and RNase free water. The ddPCR assays were performed using the QX200 EvaGreen supermix, cDNA, and pairs of primer for the actual assays (Bio-Rad, Hercules, CA, USA), as previously described ^24,25^. The amplified PCR products were quantified using Bio-Rad QX200 droplet reader and analyzed with its associated QuantaSoft software. We used 4 internal control genes (*GAPDH*, *PPIA*, *HPRT1*, and *RPLP0*)^29^ to be quantified in the samples for normalization purpose. There was no difference in the expression of the tested genes after normalization with any of the internal control genes. Thus, we decided to normalize with GAPDH for the figure purpose. Mann-Whitney’s test was used to analyze the differential expression data from ddPCR for the statistical analysis.

### Expression correlation and Network Analysis

The expression levels of differentially expressed coding RNAs and lncRNAs in all 18 corneal samples were imported into RStudio Version 0.98.1062 (RStudio, Boston, MA, USA) ^30^ from CSV raw data files. The combined coding RNAs and lncRNAs data was structured into a table where the rows were the mapped RNA names and the columns were the case or control individual deidentified with a code name. Repeated gene names were removed and genes with no expression levels in all controls and cases were also removed. The correlation between all Coding RNA and lncRNA expression levels were determined for the controls data and the cases data separately using R’s built-in Pearson correlation function (cor). Using R psych package’s test of significance for correlations (r.test), the p-value comparing the respective mapped gene’s control and case correlations were calculated. The false discovery rate (FDR) was calculated from the p-values using the R’s built-in Benjamini–Hochberg procedure function (p.adjust with “BH” parameter). We removed redundant correlations between each gene and itself as well as duplicated gene correlations (A-B is the same as B-A). Each correlation was identified along with their respective expression levels in cases and controls, p-values, and FDR.

The list of expression correlations was filtered with the criteria of an FDR ≤ 0.05 and having at least a mean expression level ≥ 20 in the cases or the controls of each gene. The entire filtered list of mapped RNAs and lncRNAs that had FDR<0.05 and had at least one mean expression level above 20 in cases or controls were visualized using Cytoscape version 3.5.1 (Cytoscape Consortium, New York, NY) ^31^ to identify gene expression networks of coding RNAs and lncRNAs. Expression network hubs were defined as network nodes with multiple correlations to other coding RNAs and/or lncRNAs. It was expected that this correlation expression analysis among coding RNAs and lncRNAs would help identify biologically important lncRNAs in human cornea. The list of correlated coding genes was loaded to WebGestalt (www.webgestalt.org) 2013 version for gene ontology enrichment and KEGG pathway analyses.^27^

## Results

### Identification of differentially expressed coding RNAs

After sequencing, each RNA sample generated 20-44 million paired reads with 100% sample index match, > 96% bases with Q30, and mean quality score > 38. Overall, 62% - 88% of sequencing reads for all RNA samples were successfully mapped to the transcriptome file from Ensembl database, containing a total of 212,368 transcript isoforms. Using a cutoff of zero transcripts, there were a total 95,954 and 103,440 expressed transcripts in all the normal controls and KC-affected cornea samples, respectively. The number of expressed transcripts in each sample ranged from 120,000 to 139,000. Using a cutoff of at least one count of normalized transcripts, the number of expressed transcripts in each sample ranged from 35,000 to 45,000. There is a total of 14,111 and 19,608 expressed transcripts (≥1 count) in all the normal controls and KC-affected corneal samples, respectively. The mean expression levels of these shared transcripts (≥ 1 count) were 142 reads in control corneas and 63 read counts in the keratoconic corneas.

Using the cutoff with |fold change| ≥ 2 & FDR ≤ 0.05, we identified 436 differentially expressed coding RNAs (159 up-regulated and 277 down-regulated) (**Supplemental Table 1**). Our analysis was consistent with previous expression studies by identifying differential expression of many known KC-related genes, including but not limited to *TGFBR3*, *TIMP1*, *FBLN1*, *TIMP3*, *AQP5*, and *SFRP1*. The top 20 up- and down-regulated coding genes, showing the highest fold changes, have been listed in **Table 2**.

**Table 2.**
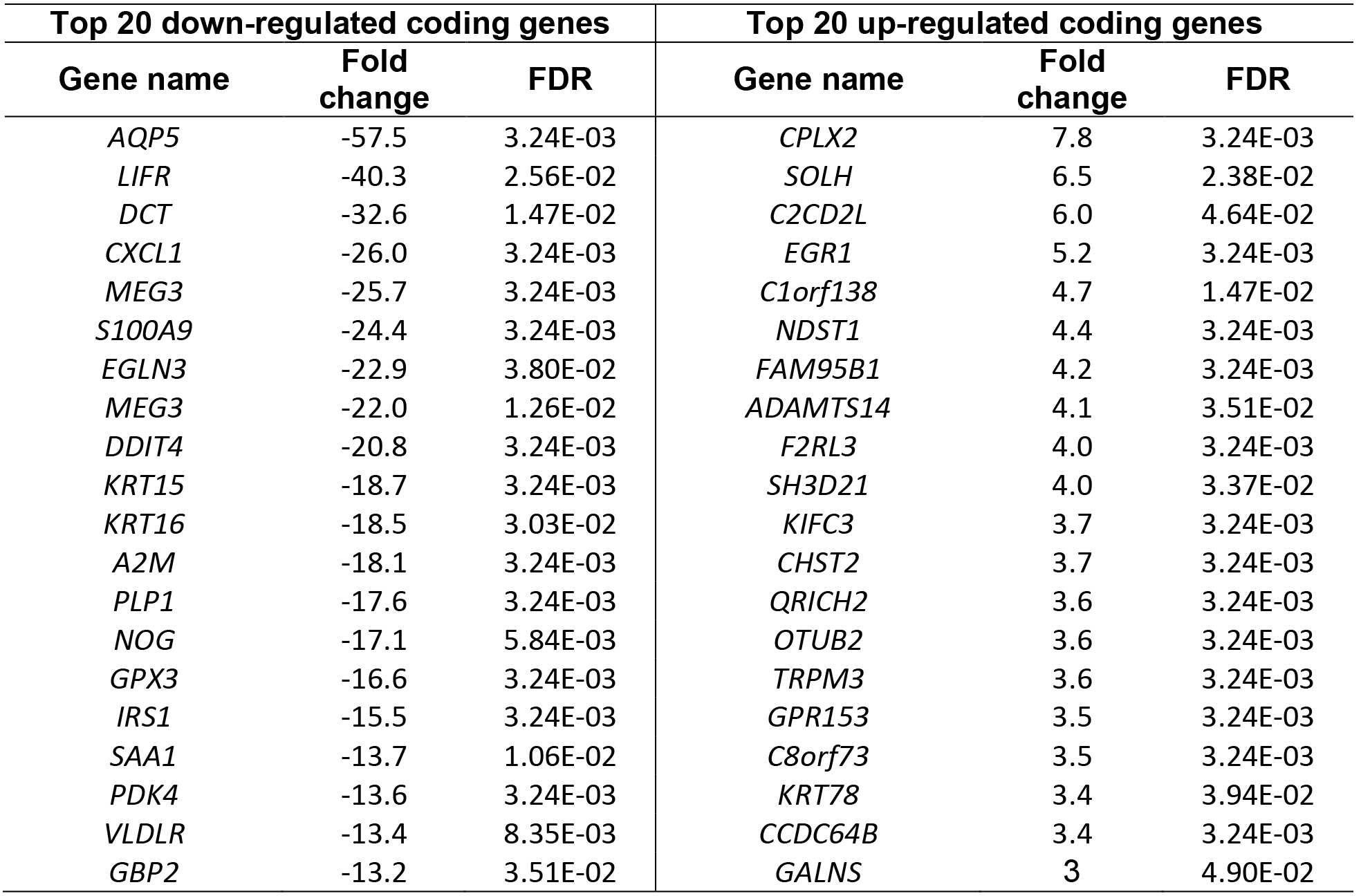
Top coding genes with significant differential expression in KC-affected corneas

Using WebGestalt with the differentially expressed coding RNAs, our gene ontology analysis revealed disruption in different biological processes (including cellular proliferation, differentiation, locomotion, migration, cellular stress and wound response), molecular functions (including glycosaminoglycan binding, iron ion binding, antioxidant activity, and insulin like growth factor binding), and cellular components (including adherens junctions, extracellular matrix region, and basement membrane) in the keratoconic corneas.

### Validation of differentially expressed coding RNAs

Out of the 436 differentially expressed coding genes, we selected 21 genes for validation by ddPCR (**Table 3**). Of these 21 genes, 19 were confirmed to have significant and similar expression trend to the RNA-Seq results (Figures 1A and **1B**). The other two genes (*APOD* and *IFI6*) demonstrated similar expression trends but did not reach statistical significance.

**Table 3.**
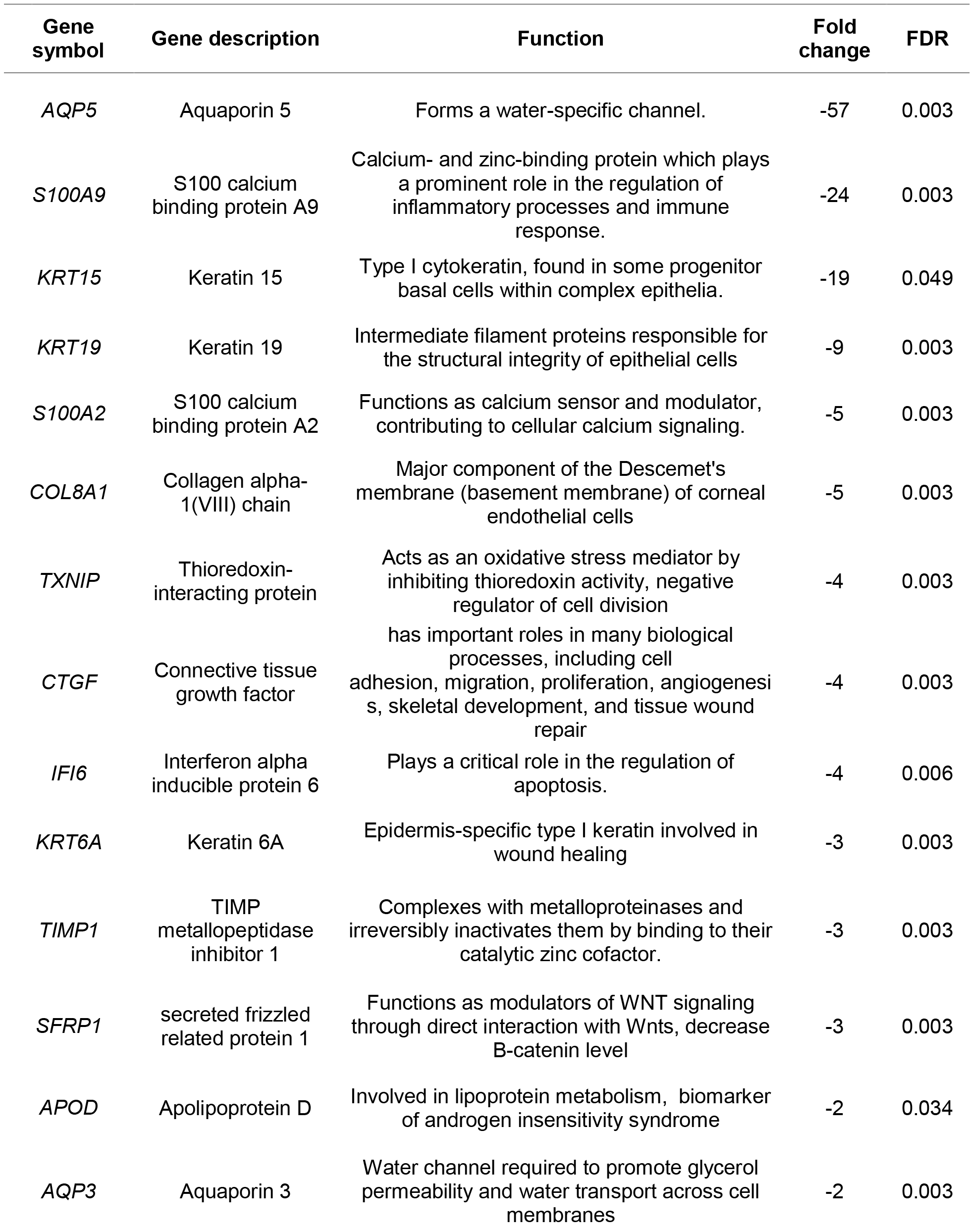
Twenty one differentially expressed coding genes to be validated using ddPCR.

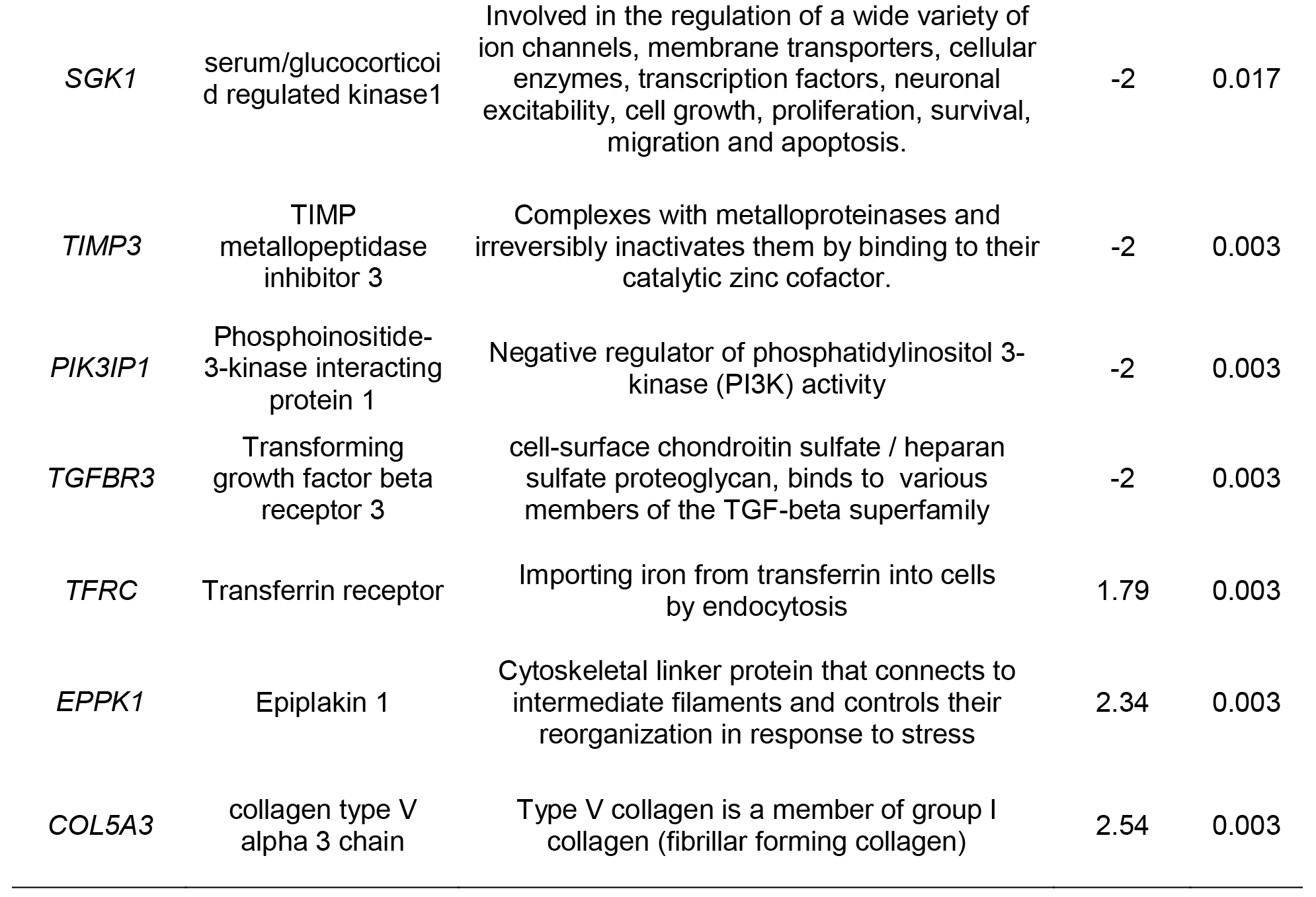

**Figure 1.**
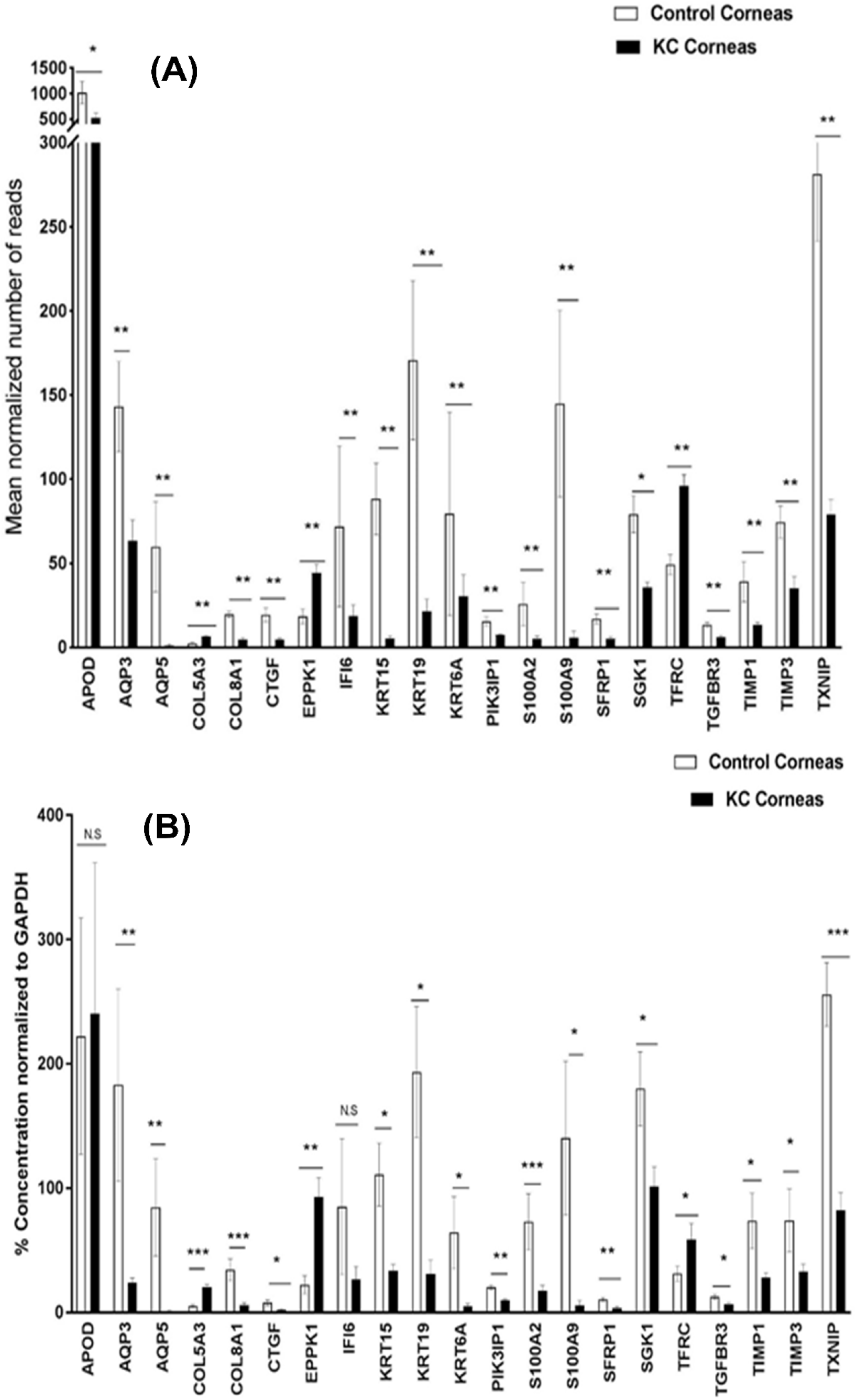
Differentially expressed coding genes in keratoconus-affected human corneas identified by RNA sequencing (A) and droplet digital PCR (ddPCR). For panel A, ** FDR < 0.01, * 0.01< FDR < 0.05. For panel B, ***p- value < 0.001, ** p-value < 0.01, * 0.01< p-value < 0.05. Error bars in panels A and B represent standard error.

### Identification of differentially expressed lncRNAs

A total of 15,966 lncRNAs were expressed in the control and keratoconic cornea samples. Using a fold change cutoff of ≥ 2 & FDR value cutoff of ≤ 0.05, we identified 586 differentially expressed lncRNAs (343 up-regulated and 243 down-regulated) (**Supplemental Table 2**). Next, differentially expressed coding RNAs and lncRNAs were used to perform a correlation analysis.

### Validation of differentially expressed IncRNAs

We selected 7 differentially expressed lncRNAs for ddPCR validation (**Table 4**). Specific probe-based ddPCR assays with FAM labeling were designed to validate the differential expression of these lncRNAs in KC-affected corneas versus the controls (**Figure 2A**). The four selected internal reference genes were also used to normalize the expression of these lncRNAs. Five lncRNA showed a similar trend to RNA-Seq results (*lnc-WNT4-2:1*, *MEG3*, *lnc-BLID-5:1*, *lnc-SCP2-2:1*, and *lnc-ALDH3A2-2:1*). However, *NONHSAT174656* and *lnc-DLG-1:1* didn’t show any significant differential expression in KC-affected corneas versus controls (**Figure 2A & 2B**).

**Table 4.**
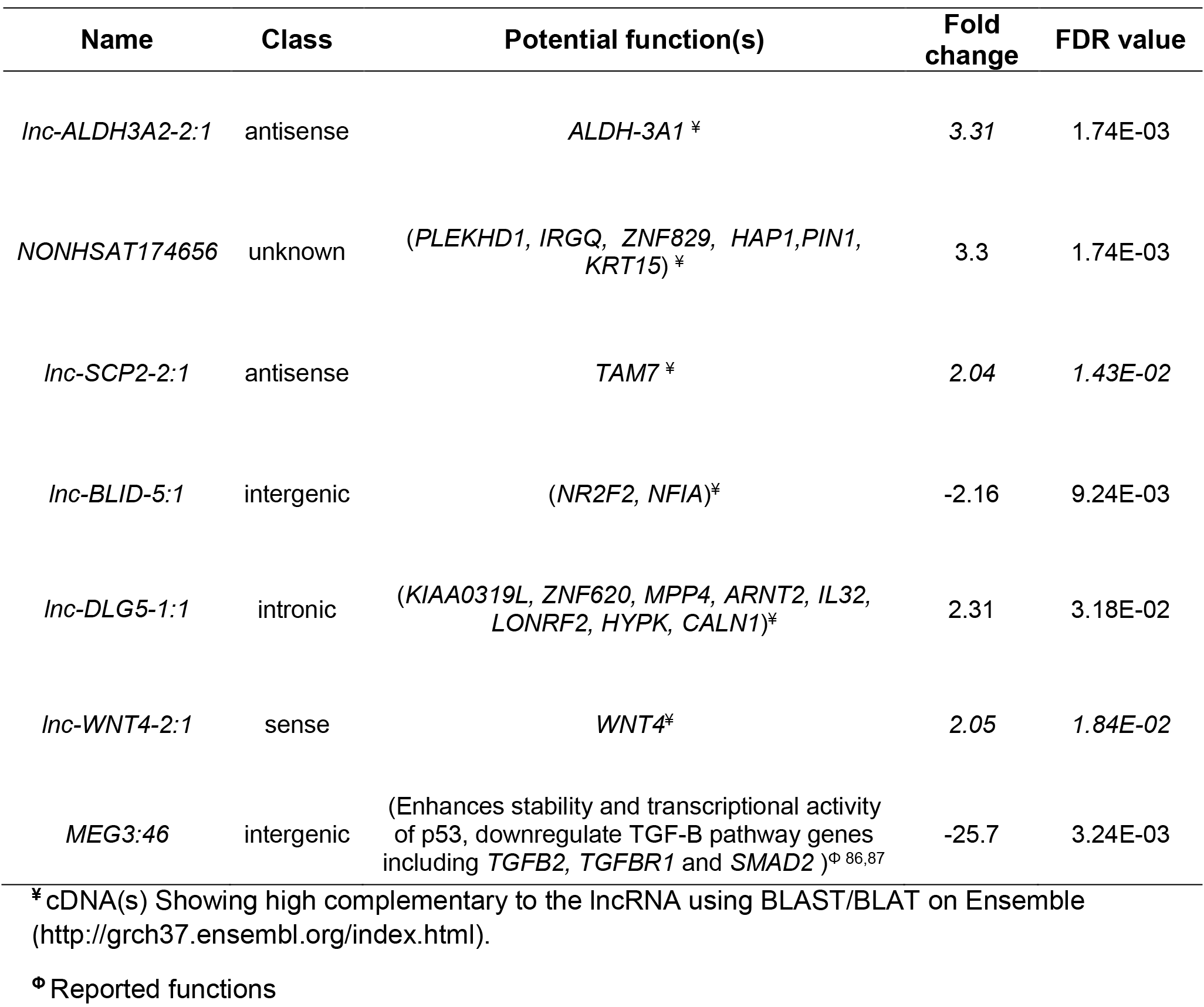
Seven differentially expressed lncRNAs to be validated using ddPCR

**Figure 2.**
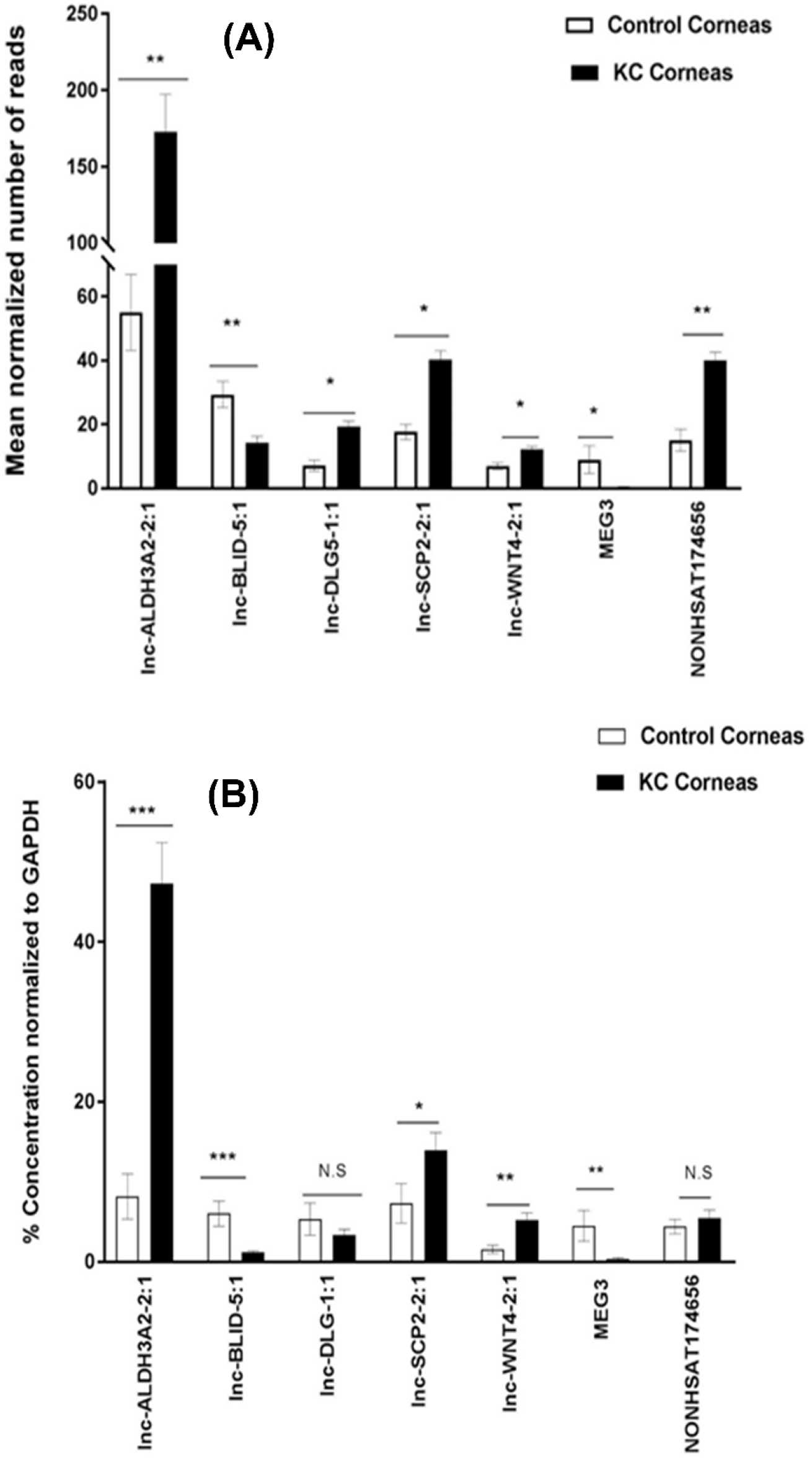
Differentially expressed long non-coding RNAs (lncRNAs) in keratoconus-affected human corneas identified using by RNA-Seq (A) and droplet digital PCR (ddPCR) (B). For panel A, ** FDR < 0.01, * 0.01 < FDR < 0.05. For panel B, *** p- value < 0.001, ** p-value < 0.01, * 0.01< p-value < 0.05. Error bars in panels A and B represent standard error.

### Correlation analysis of coding and noncoding RNAs

The expression correlation analysis was performed using the differentially expressed RNAs (436 coding RNAs and 586 lncRNAs) from 18 human corneal samples. The analysis identified a total of 492,135 pairs of RNAs with expression correlations in cases or controls. After comparing the correlations between cases and controls, a total of 352 significant differential correlations were identified with a FDR ≤ 0.05. After applying a minimum mean expression of 20 for at least one of the two paired transcripts in cases or controls, we identified 296 pairs of significant differential expression correlations between cases and controls (Supplemental Table 3). Many coding and noncoding RNAs were identified in multiple correlation pairs. These included *A2M*, *APOD*, *CCND2*, *DPYSL2*, *ELOVL5*, *GNAZ*, *LGALSL*, *MUC16*, *RAB11FIP4*, *RGS2*, *SAA1*, *SLC9A7*, *lnc-SCP2-2:1*, *lnc-DLG5-1:1*, *lnc-BLID-5:1*, *lnc-ALDH3A2-2:1*, *NONHSAT149712*, *NONHSAT154490*, *NONHSAT162589*, *NONHSAT174656*, *NONHSAT175650_1*, *NONHSAT179872*, *NONHSAT189057*, *NONHSAT193811*, *NONHSAT206355*, *NONHSAT211631*, and *NONHSAT217671*. We then used Cytoscape to visualize the whole expression network of these 296 pairs (**Figure 3**). This network analysis revealed a number of genes with significant correlations in the keratoconic corneas versus the unaffected normal controls. (**Table 5**). We identified 117 coding RNAs in the significantly differential expression correlations. These 117 genes were uploaded to WebGestalt for gene ontology analysis, and a number pathways were significantly associated, including negative regulation of cell migration and cell motility, extracellular space, cellular response to cytokine stimulus, cell surface receptor signaling pathway, regulation of cell adhesion, and epithelial tube formation.

**Table 5.** Coding and long noncoding RNAs with significant differences in expression correlation (with a reasonable corneal expressions) in KC-affected corneas versus the controls

| Gene1 | Gene2 | Correlation |  | FDR | Gene 1 mean expression |  | Gene 2 mean expression |  |
| --- | --- | --- | --- | --- | --- | --- | --- | --- |
|  |  | Control | KC |  | Control | KC | Control | KC |
| *Inc-TRIM59-2:1* | *HLA-DRA* | -0.8 | 0.8 | 2.1E-02 | 7902 | 2906 | 13 | 6 |
| *Inc-RPL13A-1:8* | *C3* | -0.9 | 0.8 | 1.9E-02 | 1863 | 848 | 11 | 1 |
| *NONHSAT176198* | *NONHSAT174656* | 0.9 | -0.7 | 2.4E-02 | 1065 | 445 | 17 | 57 |
| *NONHSAT193409* | *Inc-ALDH3A2-2:1* | -0.9 | 0.9 | 5.4E-03 | 637 | 239 | 94 | 310 |
| *NONHSAT217671* | *NONHSAT169324* | 0.9 | -0.8 | 2.6E-02 | 635 | 215 | 222 | 101 |
| *NONHSAT217671* | *Inc-ALDH3A2-2:1* | -0.8 | 0.8 | 3.5E-02 | 635 | 215 | 94 | 310 |
| *NONHSAT193408* | *Inc-ALDH3A2-2:1* | -0.8 | 0.9 | 1.2E-02 | 632 | 230 | 94 | 310 |
| *NONHSAT152360_1* | *LGALSL* | 0.8 | -0.7 | 4.9E-02 | 421 | 161 | 54 | 121 |
| *Inc-KIAA1107-2:1* | *A2M* | 0.8 | -0.7 | 4.9E-02 | 256 | 128 | 5 | 0 |
| *NONHSAT193385* | *NONHSAT175650_1* | 0.9 | -0.9 | 1.2E-02 | 252 | 125 | 4 | 16 |
| *Inc-C11orf41-1:1* | *TSC22D1* | -0.9 | 0.8 | 2.5E-02 | 225 | 103 | 78 | 17 |
| *Inc-APH1B-1:1_1* | *CLIC4* | -0.9 | 0.7 | 3.5E-02 | 170 | 28 | 13 | 6 |
| *NONHSAT189057* | *Inc-SCP2-2:1* | 0.9 | -0.8 | 5.8E-03 | 149 | 43 | 30 | 62 |
| *NONHSAT209018* | *Inc-C3orf77-2:1* | -0.8 | 0.9 | 2.0E-02 | 115 | 232 | 4 | 17 |
| *Inc-SBDS-16:1* | *STK35* | -0.8 | 0.9 | 4.0E-02 | 111 | 253 | 60 | 134 |
| *NONHSAT151930* | *Inc-DLG5-1:1_1* | -0.9 | 0.8 | 7.4E-03 | 87 | 33 | 8 | 17 |
| *NONHSAT151930* | *Inc-DLG5-1:1* | -0.8 | 0.9 | 7.9E-03 | 87 | 33 | 8 | 17 |
| *NONHSAT157212* | *Inc-SCP2-2:1* | -0.8 | 0.9 | 2.1E-02 | 71 | 148 | 30 | 62 |
| *NONHSAT157212* | *NONHSAT154490* | -0.8 | 0.7 | 3.5E-02 | 71 | 148 | 19 | 78 |
| *STK35* | *A2M* | 0.8 | -0.8 | 1.1E-02 | 60 | 134 | 5 | 0 |
| *TSPAN6* | *LGALSL* | 0.8 | -0.8 | 2.5E-02 | 56 | 19 | 54 | 121 |
| *Inc-WNT1-5:1* | *ARL4A* | 0.9 | -0.8 | 1.4E-02 | 55 | 118 | 6 | 2 |
| *NONHSAT186356* | *NONHSAT148593* | 0.9 | -0.8 | 1.3E-02 | 54 | 15 | 436 | 110 |
| *NONHSAT154490* | *Inc-RP11-770G2.3.1-2:1* | -0.8 | 0.8 | 3.8E-02 | 19 | 78 | 29 | 67 |
| *SFRP1* | *APOD* | -0.8 | 0.8 | 1.9E-02 | 17 | 5 | 1118 | 511 |
| *NONHSAT186356* | *NONHSAT148593* | 0.9 | -0.8 | 1.3E-02 | 54 | 15 | 436 | 110 |
| *Inc-FAM174B-7:2* | *SLC2A1* | -0.8 | 0.9 | 4.0E-02 | 11 | 36 | 182 | 62 |
| *NONHSAT151601* | *Inc-RP1-239B22.1.1-2:3* | -0.9 | 0.8 | 4.1E-03 | 13 | 29 | 61 | 28 |
| *NONHSAT152805* | *LGALSL* | -0.8 | 0.8 | 2.6E-02 | 8 | 19 | 54 | 121 |
| *CLIC4* | *CA12_1* | -0.9 | 0.8 | 2.3E-02 | 13 | 6 | 34 | 6 |

**Figure 3.**
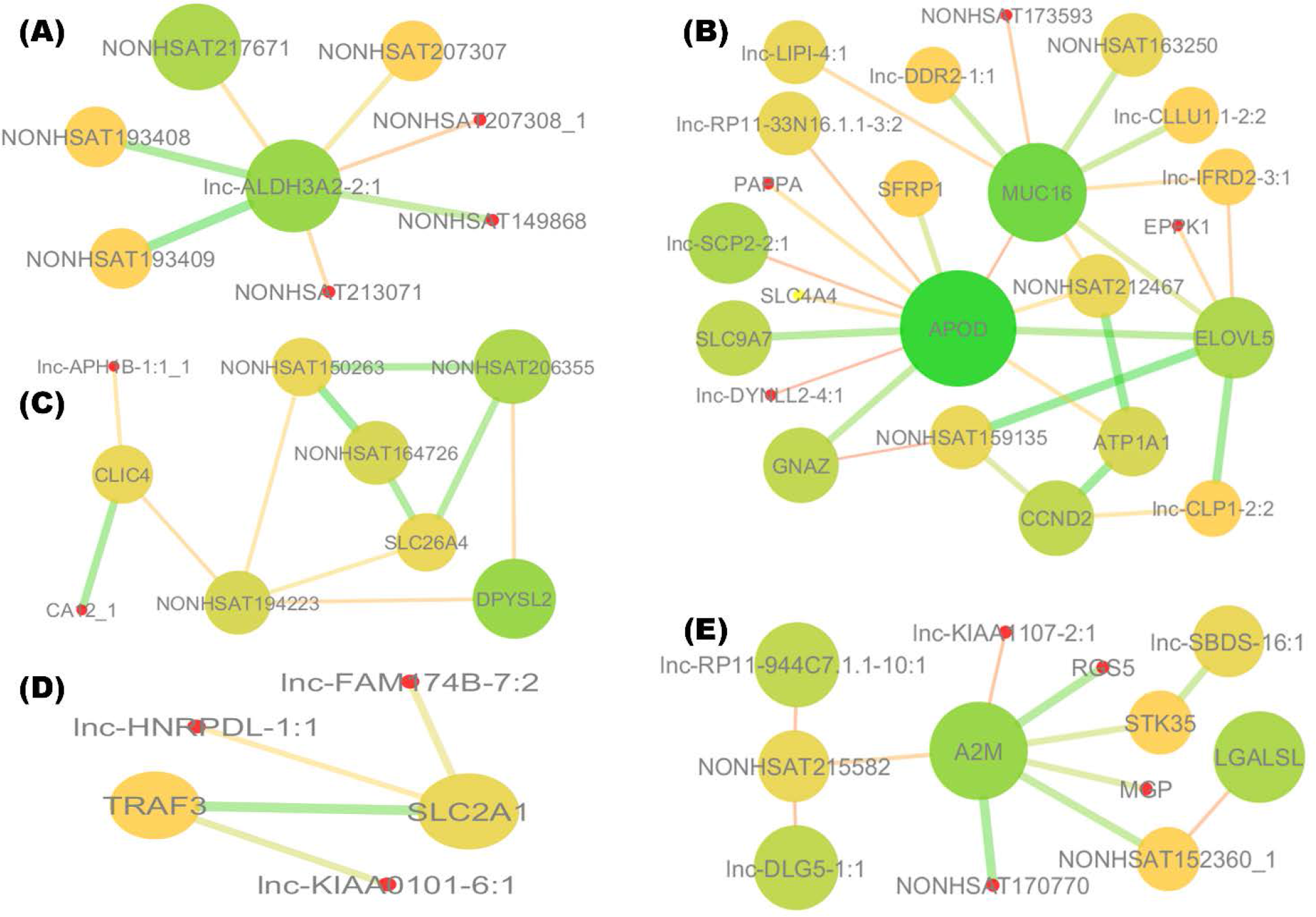
Correlation networks of the differentially expressed coding RNAs and lncRNAs displayed by Cytoscape. The nodes represent mapped RNAs colored and sized by the degree of correlations with other RNAs. The edges are sized and colored based on the significant differences (i.e., false discovery rate, FDR) between controls and cases. Correlation networks were shown for the *Lnc-ALDH3A2 −2:1* (A), *APOD* (B), *CLIC4* (C), *SLC2A1* (D), and *A2M* (E) genes.

## Discussion

We have successfully used Total RNA-Seq to identify the differentially expressed coding RNAs and lncRNAs in KC-affected whole corneal tissues. WebGestalt-based gene ontology analysis for the 436 differentially expressed coding RNAs revealed disturbances in various corneal functions including cellular adhesion, oxidative stress response, proliferation, migration, apoptosis, and wound healing. We also identified significant expression differences in non-coding RNAs in KC corneas. These data were used to implicate many potential network hubs that may play potential roles in KC pathogenesis.

KC is a complex disorder with both genetic and environmental factors. Epithelial and stromal layers of the cornea show the prominent histopathological changes during KC progression. A better understanding of the key molecular players involved in KC-associated corneal pathogenesis may provide potential therapeutic targets for disease treatment. Previous studies used either the whole corneal tissues, corneal epithelial cells, stromal fibroblasts, or even tears to examine the differentially expressed coding genes or proteins in KC ^18–20,22,23,32^. Researchers used various techniques in these studies including RNA-Seq, microarray, RT-PCR, mass spectrometry, and western blots ^17,18,22,32–34^. Recently, Kabza et al. have used RNA-Seq for KC-affected corneas; however, this study was complicated by the use of other diseased corneas as controls. Interestingly our data shows similarity in the differentially expressed coding genes with this recent RNA-Seq study (35 of the up-regulated genes including: *COL5A3*, *COL28A1*, *IL6R*, *TFRC*, and 135 of the down-regulated genes including: *TIMP1*, *TIMP3*, *IFI6*, *IRS1*, *IGFBP2*, *DDIT4*)^32^ (**Supplemental Table 4**).

We identified a significant down regulation in the cytokeratin (*KRT6A*, *KRT13*, *KRT15* and *KRT19*) and collagen (*COL8A1* and *COL8A2*) genes, consistent with the compromised integrity of the KC corneal tissues. Type VIII collagens are normally expressed in the Descemet’s membrane of human cornea^35^ and targeted inactivation of Col8a1 and Col8a2 has led to corneal abnormalities in mice^36^. Type V collagens are normally expressed in corneal stroma and Bowman’s layer ^37^. Radda et al. have not shown expression changes in type V collagen in KC cornea^38^; however, our sequencing and validation data shows upregulation of *COL5A3* in the KC-affected cornea. These contradictory results could be due to the differences in corneal sample processing (collagen types were separated using salt fractionation and thermal gelation in the pepsin-soluble fraction of the lyophilized normal and KC human corneal tissues)^38^.

Our study confirms the potential role of oxidative stress in KC-affected corneas. This is consistent with previous studies, suggesting the involvement of cellular stress in KC pathology^39,40^. We have identified differential expression of many genes involved in response to oxidative stress including *GSTO1*, *SGK1*, *APOD*, *KLF9*, *GPX3*, *STK39*, *TXNIP*, and *lnc-ALDH3A2-2:1*. The majority of the coding genes involved in defense mechanism for oxidative stress were down-regulated. However, *lnc-ALDH3A2-2:1* is found to have more than 3-fold increased expression and correlated to other lncRNAs (Figure 3A). *Lnc-ALDH3A2-2:1*, according to Lncipedia database^41^, is classified as an antisense transcript which overlaps with the protein coding gene *ALDH3A2* on the opposite strand. However, using BLASTN, we found that *lnc-ALDH3A2-2:1* sequence overlaps with the last exon of *ALDH3A1* gene. The *ALDH3A1* transcript did not show significant differential expression in our study; however, lncRNAs may alter the protein expression level without affecting the coding RNA level^42^. For example, some lncRNAs can control expression of coding genes by acting as molecular decoys (binds and titrates away RNA-binding proteins such as transcription factors, chromatin modifiers, or other regulatory factors). Another factor that can influence the impact of oxidative stress is disruption of iron homeostasis, which can lead to various corneal diseases including KC^20,21,43,44^. Fleisher ring, a ring of iron deposition in the corneal cone, is a typical clinical sign for KC^45^. Importantly, TFRC, a cellular receptor for iron-ferritin complex uptake^46^, has been identified to have elevated expression in KC cornea with both RNA-Seq and ddPCR in our study. This elevated expression in KC-affected corneas may result from iron deficiency^47^. We have also found down-regulation of the *FTL* gene (Ferritin Light Chain) in KC corneas. Significantly, mutations in *FTL* have been linked to iron accumulation in the brain, causing neurodegenerative disease^48^.

We observed differential expression of many genes involved in corneal wound healing such as *EPPK1*, *AQP3*, *AQP5*, *SLC2A1* and *CTGF*. Corneal wound healing is a complex process controlled by the integrated actions of multiple growth factors, cytokines, and proteases produced by epithelial cells, stromal keratocytes, and inflammatory cells ^49^. However, unregulated healing processes, due to genetic or epigenetic factors, can lead to impairment in cellular function ^50,51^. Mace et al. have shown anti-proliferative and hyper-apoptotic phenotype in KC-affected cornea which also reveal a defect in wound healing process^33^. CTGF is expressed in multiple ocular tissues as a downstream signaling molecule in the TGF-β pathway^52^. Importantly, loss of *CTGF* impairs efficient corneal wound healing^53^. We have found that *CTGF* was significantly down-regulated in KC-affected corneas, which is consistent with previous studies ^24,32^. *EPPK1* encodes a cytoskeletal linker protein called epiplakin, which maintains the integrity of intermediate filaments networks in epithelial cells ^54,55^. Migration-dependent wound healing improvement has been found in epiplakin-null epithelium ^56^. Our data shows an up-regulation of *EPPK1*, suggesting the involvement of *EPPK1* in the impaired corneal wound healing in KC. Interestingly, SLC2A1 (solute carrier family 2 member 1, also called glucose transporter-1) is up-regulated during corneal epithelium wound healing^57^. In our results, *SLC2A1* shows a 3-fold down-regulation in KC-affected corneas, which may reflect a lesser glucose uptake. This is intriguing because there appears to be a negative association between diabetes mellitus and KC (i.e. a reduced risk of developing KC in diabetic patients)^58–61^. Moreover, it has been reported that diabetic patients have higher corneal thickness than normal individuals^57^.

Aquaporins (AQPs) are a ubiquitous family of transmembrane water channels that are important to maintain tear film osmolarity, stromal layer thickness, and corneal transparency ^62^. *AQP3* and *AQP5* are highly expressed in the cornea and important for corneal wound healing ^62^. Both AQPs show reduced expressions in the KC-affected corneas. AQP3 channels are located in the epithelial cells. *Aqp3* null mice demonstrate significant impairment in corneal re-epithelialization and delayed wound healing ^63^. *AQP5* transcript absence in KC-affected corneas have been reported by Rabinowitz et al. ^64^, which is consistent with our results (**Table 2**). Although the lack of *AQP5* differential expression has been reported in KC-affected corneas by Garfias et al., their controls were heterogeneous (normal corneal epithelium, sclerocorneal rims, and healthy corneal buttons) which may explain the controversial results^65^. *Aqp5* knockout mice are reported to have hyperosmolar tears ^66^. Hyperosmolarity is considered a biomarker for dry eyes in human ^67^. Dry eye symptoms have been reported in KC patients^68–70^, thus reduced expression of *AQP5* might account for dry eye symptoms in some KC patients. Extracellular matrix (ECM) regulatory pathways are important for corneal integrity; however dysregulation in these pathways have been reported with KC progression ^24,45^. Previous studies have shown an up-regulation of ECM degradative enzymes (such as matrix metalloproteinases and cathepsin) and/or down-regulation of their inhibitors (such as α1-protease inhibitor, α2-macroglobulin, secretory Leukocyte Peptidase Inhibitor, and TIMPs) ^32,39,71^. Our data shows significant down-regulation in proteinases inhibitors (*TIMP-1*, *TIMP-3*, and *A2M*, and *SLPI*), but interestingly no significant change with ECM proteinases.

WNT signaling pathway is essential for normal corneal development and a missense coding variant (rs121908120, c.1145T>A, p.228Phe>Ile) in the *WNT10A* gene has been associated with KC risk^72^. Moreover, using RNA-Seq, a recent study showed dysregulation in WNT signaling in the KC-affected corneal epithelium.^34^ We found some evidence of WNT signaling disruption represented by downregulation of *ANGPTL7*, *TGFBR3*, *SFRP1* and upregulation of *FZD8* (frizzled class receptor 8) and *EGR1* (early growth response 1). *ANGPTL7* encodes angiopoietin-like 7 and is abundantly expressed in keratocytes to maintain corneal avascularity and transparency^73^. The SFRP1 protein belongs to glycoproteins family which inhibits WNT signaling by preventing the formation of the WNT-FZD complex. Iqbal et al. have shown an increased expression of SFRP1 in the KC-affected corneal epithelium^74^; however, in tears from KC patients, SFRP1 expression was reduced in KC ^75^. *TGFBR3* has been found to be a suppressor to WNT signaling through binding to Wnt3a and β-catenin ^76^. We observed that *Lnc-WNT4-2:1* was up-regulated more than two-fold in KC versus controls. *Lnc-WNT4-2:1* is classified as a sense transcript which overlaps with protein coding gene on the same strand. Interestingly, CPAT (Coding-Potential Assessment Tool using an alignment-free logistic regression model) predicts a coding potential for this lncRNA (https://hg19.lncipedia.org/)^41^. BLASTN analysis indicates *lnc-WNT4-2:1* overlaps with exon 5 of the *WNT4* gene. Although our data did not show any significant change in *WNT4* transcript, this does not rule out the potential role of *lnc-WNT4-2:1* in regulating WNT signaling pathway.

Another pathway we hypothesize to play an important role in KC pathogenesis is the PI3K/AKT pathway. This pathway is involved in cell cycle regulation and cell proliferation in the cornea^77^. Our data shows a down-regulation of negative regulators of PI3K/AKT pathway, including *PIK3IP1*, *DDIT4*, *IGFBP-3*, -*4*, and *-5. PIK3IP1* has been suggested to suppress AKT activity by inhibiting the expression of PI3K ^78^. PIK3IP1 has previously been reported to be down-regulated in KC-affected corneas^32^. IGFBPs belong to a family of six structurally similar proteins which antagonize the binding of insulin growth factors to their receptors, thus, controlling cell survival, mitogenesis, and differentiation ^79^. Our results indicate that IGFBP-3,-4, and-5 have 10-fold, 3-fold, and 8-fold reduced expression in KC corneas, respectively. Our findings are consistent with previous studies that have reported significantly reduced expression of IGFBP3 and IGFBP5 by 11- and 14-fold in keratocytes isolated from KC-affected corneas compared to normal corneas ^17^.

We have combined data from the differentially expressed coding RNAs and lncRNA to illustrate an integrated expression correlation network using cytoscape. For the first time, we have identified pairs of RNAs showing significantly different correlations in KC patients vs. controls (**Table 5**). Functions of most of the identified genes are still not clear, especially in the cornea. *APOD* and *SFRP1* showed significant positive correlation in KC, with both being down-regulated (**Figure 3B**). Both genes were reported to have tumor-suppressive functions including DNA-repair, induction of apoptosis, detoxification, differentiation, and transcriptional regulation ^80^. Apolipoprotein D (APOD) is a stroma specific protein that acts as lipophilic molecules carrier^81^. SFRP1 is a negative regulator of WNT signaling, as we mentioned earlier, but it is unclear whether WNT signaling disruption is a cause for the inverse correlation between the two genes or not.

Moreover, *CLIC4* and *CA12-1* are two other genes that showed positive correlation, both being down-regulated in KC (**Figure 3C**). CLIC4 is a chloride intracellular channel, and was found to be important for cellular adhesion and wound healing ^82,83^. Carbonic anhydrase 12 (CA12) is a membrane bound enzyme which has been upregulated in glaucoma^84^. We suggest that positive correlation between the two genes is related to corneal edema and loss of transparency associated with KC. Corneal stromal swelling have been reported after addition of carbonic anhydrase inhibitors or removal of bicarbonate anion, due to inhibition of endothelium ion active pump. Chloride transport was also found to play a role in maintaining stromal hydration level ^85^.

Despite of the significant findings, our study does have several limitations. First is the relatively small sample size of cases and controls. It will be necessary to replicate our findings in additional datasets with KC-affected and normal human corneas. Second, age, gender, and ethnicity were not matched completely between cases and controls. Third, our expression study is limited by transcript quantification only. Additional proteomics or protein-based studies will be necessary to confirm the differential gene expression in KC-affected corneas.

In summary, we have performed a comprehensive differential expression profiling of both coding and long noncoding RNAs in KC-affected human cornea samples in comparison to normal unaffected human cornea samples. We have not only confirmed many previous reported KC-related differential expressed genes, but also identified a number of novel coding and noncoding RNAs involved in KC pathogenesis. Our expression correlation analysis between differentially expressed coding RNAs and lncRNAs has identified a number of potential expression regulators and provided potential functional annotation for lncRNAs with unknown functions.

## Acknowledgements

The authors highly appreciate all the donated human corneal tissue samples from all individual donors. Without such samples, this study would not be possible.

